# Intra-Articular Sprouting Of Nociceptors Accompanies Progressive Osteoarthritis: Comparative Evidence In Four Murine Models

**DOI:** 10.1101/2023.06.30.547216

**Authors:** Alia M. Obeidat, Shingo Ishihara, Jun Li, Lindsey Lammlin, Lucas Junginger, Tristan Maerz, Richard J. Miller, Rachel E. Miller, Anne-Marie Malfait

## Abstract

**Objective:** Knee joints are densely innervated by nociceptors. Sprouting of nociceptors has been reported in late-stage osteoarthritis (OA), both in human knees and in rodent models. Here, we sought to describe progressive nociceptor remodeling in four mouse models of knee OA, capturing early and late-stage disease.

**Methods:** Sham surgery, destabilization of the medial meniscus (DMM), partial meniscectomy (PMX), or non-invasive anterior cruciate ligament rupture (ACLR) was performed in the right knee of 10-12-week old male C57BL/6 NaV1.8-tdTomato mice. Mice were euthanized (1) 4, 8 or 16 weeks after DMM or sham surgery; (2) 4 or 12 weeks after PMX or sham; (3) 1 or 4 weeks after ACLR injury or sham. Additionally, a cohort of naïve male wildtype mice was evaluated at 6 and 24 months. Twenty-μm thick mid-joint cryosections were assessed qualitatively and quantitatively for NaV1.8+ and PGP9.5+ innervation. Cartilage damage (using a modified OARSI score), synovitis, and osteophytes were assessed blindly.

**Results:** Progressive OA developed in the medial compartment after DMM, PMX, and ACLR. Synovitis and associated neo-innervation by nociceptors peaked in early-stage OA. In the subchondral bone, channels containing sprouting nociceptors appeared early, and progressed with worsening joint damage. Two-year old mice developed primary OA in both the medial and the lateral compartment, accompanied with neuroplasticity in the synovium and the subchondral bone. All 4 models had an increased nerve signal in osteophytes.

**Conclusion:** Anatomical neuroplasticity of nociceptors was observed in association with joint damage in 4 distinct mouse models, suggesting that it is intrinsic to OA pathology.

## INTRODUCTION

Osteoarthritis (OA), the most common form of arthritis affecting synovial joints, manifests as progressive molecular and anatomical derangements in all joint tissues, causing the joint to fail structurally and functionally (1). As a disease of the whole joint as an organ, OA is characterized by progressive pathological changes in articular cartilage, synovium, menisci, and subchondral bone, as well as in peri-articular tissues such as muscles and ligaments (2). Cellular and molecular changes in the individual tissues and their interactions have been increasingly well documented in human joints and experimental models, paving the way for identifying new targets for development of disease-modifying OA drugs (DMOADs) (3). The main symptom of OA, and the primary reason why affected individuals seek medical care is chronic pain (4,5). However, available pain management options are often not sufficiently efficacious or safe, and the wide availability of opioids has contributed to the ongoing opioid crisis (6). Hence, OA pain is a formidable worldwide problem for which safe and effective pharmacological treatments are urgently needed. This pressing medical need has resulted in an increasingly intense focus on research aimed at unraveling the mechanisms underlying pain in OA (7,8). Furthermore, our understanding of the bidirectional relationship between joint tissue damage and OA pain is limited, as exemplified by recent clinical trials with the monoclonal antibody tanezumab, which neutralizes nerve growth factor (NGF), when it was observed that successful pain relief was accompanied by rapidly progressive OA in a subset of knee OA patients. This unexplained side effect ultimately led to a halt in the development of these antibodies (9).

A critical gap that hinders the development of novel, effective pain therapeutics for OA is our incomplete understanding of the nociceptive innervation of the joint. The healthy knee joint is supplied by a complex system of sympathetic and sensory neurons, predominantly nociceptors (10–12). Nociceptors, mostly unmyelinated C-fibers, innervate all joint tissues except for cartilage, which is aneural (12). Few studies have attempted to systematically describe the precise anatomical location of nociceptor nerve endings in the knee (11,12). Importantly, a major observation emerging from recent research in human and rodent knees is that the sensory innervation of the knee is not static. In fact, it appears that the nociceptive innervation of the pathological knee joint undergoes profound alterations. For example, in human osteoarthritic knee joints and in rodent models of knee OA, calcitonin gene related peptide (CGRP)-immunopositive neurons have been reported in osteochondral channels in the sclerotic subchondral bone (13–15).

Similarly, in the late stages of experimental OA induced by surgical destabilization of the medial meniscus (DMM), nociceptor sprouts were observed in particular locations in the medial compartment, including the synovium, the meniscus, and within channels in the subchondral bone, (16). Conflicting findings have been reported in the synovium, however, with either increased nociceptive innervation in surgical models (15,16), a transient or permanent reduction of innervation in collagenase induced OA (17,18), or no changes in mechanically loaded murine joints (19). This may be attributable to varying inflammatory components in different models, as well as lack of data covering different stages of joint damage. Indeed, most published studies analyzed joints with end-stage OA, including in human knees, which are often collected at time of total joint replacement (13,20,21).

Since information on the precise anatomical distribution of free nerve endings in experimental knee OA remains limited, we sought to provide a detailed, longitudinal and systematic description of the nociceptive innervation of the mouse knee in four widely used murine OA models in order to assess whether particular changes in joint innervation were common across models. We included two surgical models - DMM and partial meniscectomy, PMX (22) -, a non-invasive model of post-traumatic OA (PTOA) induced by anterior cruciate ligament rupture (ACLR) (23), and primary age-associated OA (24,25). We selected timepoints spanning both early and later stages of disease severity for each model, as assessed by histological joint damage assessment. Furthermore, we and others have extensively documented time-dependent pain-related behaviors in each of these models (22,25–28). Here, we describe the precise distribution of intra-articular nociceptor endings at early and late disease stages, as they relate to pathological changes in the joint tissues in these 4 distinct mouse models. We found that structural neuroplasticity of joint nociceptors occurred in all four models, starting in the early stages of disease, and with a similar anatomical distribution. We conclude that nociceptor plasticity is a widely observed phenomenon in OA affected knee joints.

## MATERIALS AND METHODS

### Animals

All experiments were performed in male mice C57BL/6 (n=69), either wild-type (WT) mice (inbred at Rush) or NaV1.8Cre-tdTomato reporter mice (a gift from Dr. John Wood, University College London, London, UK) which express a bright red fluorescent tdTomato reporter in all neurons that express the voltage-gated sodium channel, NaV1.8. This neuronal subset comprises approximately 75% of dorsal root ganglion (DRG) sensory neurons, including >90% of C-fibers, as well as a fraction of Aδ-nociceptors (29,30). All animals were bred in-house, and randomly assigned to an OA model and groups. Animals were housed with food and water *ad libitum* and kept on 12-hour light cycles. Animal procedures were approved by the Institutional Animal Care and Use Committee at Rush University Medical Center or the University of Michigan. Fig. 1 shows a schematic overview of the study design.

**Figure 1:**
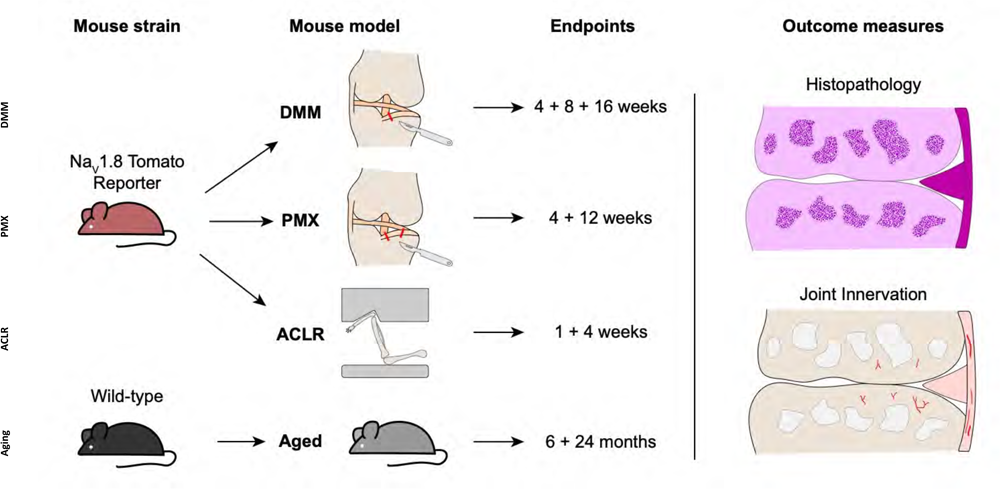
Schematic overview of male mouse strains, osteoarthritis models, timepoints, and outcome measures used in the study.

### Destabilization of the medial meniscus (DMM)

DMM (n=15) or sham surgery (n=10) was performed in the right knee of 10-week old male NaV1.8-tdTomato reporter mice (31). All DMM and sham surgeries were performed by the same surgeon (SI). Mice were sacrificed 4, 8 or 16 weeks after surgery (n=5 per group). For a detailed description of the surgeries, please see Suppl. Methods.

### Partial meniscectomy (PMX)

PMX (n=10) or sham surgery (n= 10) was performed in the right knee of 10-week old male NaV1.8-tdTomato mice, as described (22). All PMX and sham surgeries were performed by the same surgeon (JL). Mice were sacrificed 4 or 12 weeks after surgery (n=5 per group). For a complete description of PMX surgery, see Suppl. Methods.

### Anterior cruciate ligament rupture (ACLR)

At the age of 12-14 weeks, male NaV1.8-tdTomato reporter mice were randomized to a Sham group (n=5) (anesthesia and analgesia only, no injury loading) or tibial compression-based, noninvasive ACLR (n=9), using computer-aided block randomization. Within the ACLR group, mice were further randomized to two study timepoints, 1 week (n=5) or 4 weeks (n=4). These timepoints capture pre-PTOA synovitis (1 week) and established PTOA (4 weeks) (26). For a detailed description of ACLR injury, please see Suppl. Methods.

### Age-associated primary OA

Male WT C57BL/6 mice were bred in-house and aged without any interventions. Mice were sacrificed at age 26 weeks (n=5) or 2 years (n= 5), and right knees were collected.

### Knee histology and immunohistochemistry

Mice were transcardially perfused, and right knees were collected. Knees were decalcified in 14% EDTA for 2-3 weeks and cryopreserved using 30% sucrose. Twenty µm-thick coronal sections were then collected from the mid-joint area as previously described (16). Knee sections from WT mice were stained for the pan-neuronal marker, protein gene product 9.5 (PGP9.5), as described (16). Sections were blocked and incubated overnight with a primary antibody against PGP9.5 (rabbit polyclonal antibody, Sigma-Aldrich, SAB4503057; 1:100) at 4 °C, followed by a secondary antibody, anti-rabbit conjugated Alexa Fluor 633 (Molecular Probes, 1:500). Control sections were treated the same but without primary antibody. Sections were imaged using a laser-scanning confocal microscope (Olympus IX70). Images were then processed using ImageJ and Fluoview software (FV10-ASW 4.2 Viewer). Adjustments were made to brightness and contrast to reflect true colors (16). All images were treated the same in terms of adjustments to brightness and contrast to minimize bias.

### Quantification of the neuronal signal per region

All quantifications were performed by an observer blinded to the treatment groups.

In the medial and lateral synovium, NaV1.8+ and PGP9.5+ signals were quantified in mid-joint coronal sections using ImageJ. Briefly, regions of interest (ROI) were manually outlined for each section using anatomical landmarks. Thresholds were adjusted for all images similarly to control for background. The area of positive signal within each ROI was measured and was normalized to the total area of the ROI; the percentage of positive signal per ROI is reported.

NaV1.8+ and PGP9.5+ subchondral bone channels were quantified as follows: two sections (80 μm apart) were used to count the number of positive channels. The average number of positive channels was reported as previously described (16). The length of positive channels was measured using ImageJ neuroanatomy plugin (32) as follows: The lengths of all NaV1.8+ and PGP9.5+ channels per knee section were measured and the length of the longest channel was reported for that section. The distance from the tidemark was assessed by drawing a straight vertical line between the distal end of NaV1.8 and PGP9.5 positive channels and the tidemark (the proximal end being the end connected or close to the bone marrow cavity). This distance was then measured using ImageJ and the shortest distance was reported for each knee section. Since most of the age-matched sham and naïve controls lack these channels, we averaged the distances for all existing NaV1.8 and PGP9.5 positive channels in naïve and sham control knees and compared it in all models. The number of branching points was counted using a knee section with maximum branching, as shown in Suppl. Fig. 1.

### Knee histopathology

Knee sections from all groups were stained with hematoxylin (1x-Sigma HHS160) and eosin (1%-Sigma E4382) (H&E). Knee sections were evaluated for cartilage degeneration using a modified OARSI score, as described (16) (details in Suppl. Methods). OARSl scoring was performed twice by an observer blinded to treatment groups and the average of both scores was reported. For osteophyte scoring, one section with the major osteophyte was used to assess osteophyte width and maturity, as described (25,33). Osteophyte measurements were performed using Osteomeasure software (OsteoMetrics) (25,34). Synovial hyperplasia, cellularity and fibrosis were evaluated in four joint spaces (lateral femoral, medial femoral, lateral tibial, and medial tibial separately) as described (25). All scores were performed by an observer blinded to treatment groups.

### Statistical Analysis

Sample size was determined based on our previous data comparing innervation changes between sham and DMM. Ordinary two-way ANOVA was used for comparison of multiple groups and time points followed by Tukey’s multiple comparison test. For comparison in aging mice, unpaired two-tailed Student’s t test was used for pairwise comparisons. For comparison of branching between two timepoints, a Mann Whitney test was used. Statistical analyses were performed using GraphPad Prism 9. All data are presented as mean±95%CI.

## RESULTS

### Knee histopathology

We first performed a detailed histopathological analysis of the knee joints in the four models, assessing OARSI score, synovitis, and osteophyte width and maturity at different stages of disease. We confirmed that mice in all models developed cartilage degeneration (Suppl. Fig. 2A,D,G,J), synovial pathology (Suppl. Fig. 2B,E,H,K) and (Suppl. Fig. 4), and osteophytes (Suppl. Fig. 2C,F,I,L). Representative histological images of the medial side are shown in Suppl. Fig. 3. Representative images of the whole joint are shown in Suppl. Fig. 5, in addition to total joint cartilage degeneration scores and total joint synovial scores. For a detailed description of histopathological analyses performed, see Suppl. Results.

### Nociceptor sprouting

Having confirmed that mice developed progressive osteoarthritic changes in all four models, in concordance with published findings (25,27,33,35), we documented the distribution of free nerve endings in different joint tissues at all timepoints.

#### Nociceptors sprout in the synovium

We have previously reported profound neuroplasticity of intra-articular nociceptors in the medial compartment of the murine knee joint 16 weeks after DMM, where we observed sprouting of nociceptors in the medial synovium (16). Here, we assessed NaV1.8-innervation at earlier timepoints after DMM *vs.* sham surgery, and found that 4 and 8 weeks after DMM, innervation of the deeper sublining layers of the medial synovium was increased compared to sham controls (Fig. 2A-D). This occurred as early as 4 weeks after DMM, at which time synovitis peaked (Suppl. Fig. 2B), with no further increase observed by weeks 8 and 16 (Fig. 2A). Representative sections at week 4 are shown in (Fig. 2B-D), and synovial innervation 8 and 16 weeks after DMM is shown in Suppl. Fig. 6A,B,E,F. In the lateral synovium, NaV1.8-innervation decreased with age in both the DMM and sham groups starting at the 8-week timepoint (*i.e.,* 18 weeks of age), and no further decrease was observed at week 16 (Suppl. Fig. 7A). This age-related decrease of nociceptor density in the lateral synovium is compatible with our previously reported findings (16). No neo-innervation was observed in the lateral synovium at any time, concordant with the lack of synovitis or OA changes in that compartment.

**Figure 2:**
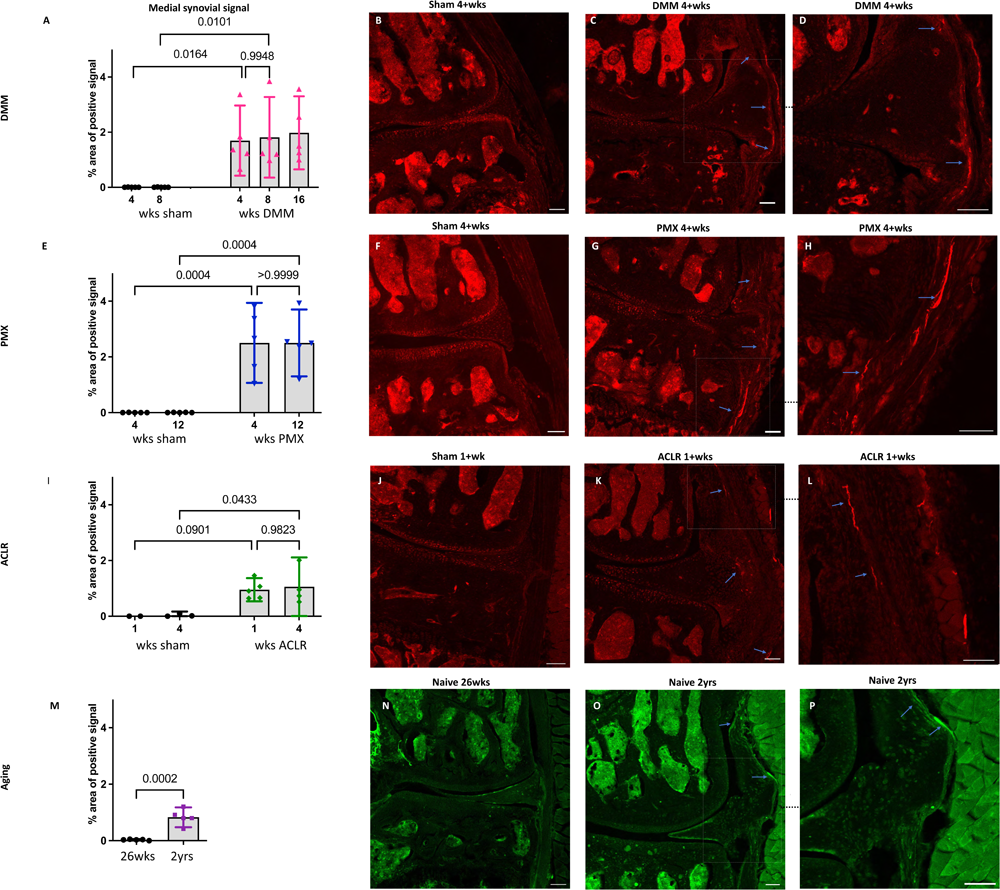
(A) Quantification of neuronal signal in the medial synovium 4, 8 and 16 weeks after sham and DMM surgeries; (B-D) Representative confocal images of NaV1.8-tdTomato mouse knees showing the nerve fibers in the medial synovium (blue arrows) at 4 weeks after sham and DMM surgery, with (D) showing a magnified image of (C) n=5/group; (E) Quantification of neuronal signal in the medial synovium 4 and 12 weeks after sham and PMX surgeries; (F-H) 4 weeks after sham, 4 weeks after PMX, and zoomed in image, respectively, n=5/group; (I) Quantification of neuronal signal in the medial synovium 1 and 4 weeks after sham and ACLR injury; (J-L) 1 week after sham, 1 week after ACLR injury, and zoomed in image, respectively, n=4-5/group; (M) Quantification of neuronal signal in the medial synovium of 26-week old and 2-year old naïve mice; (N-P) PGP9.5 staining in a 26-week old mouse, a 2-year old mouse, and zoomed in image, respectively, n=5/group. Mean ± 95% CI. Scale bar = 100 μm.

We then assessed the nociceptive innervation of the synovium in 3 additional models. For the PMX model, we selected week 4 as early disease and week 12 as late-stage disease, based on reports in the literature, and confirmed by the histological findings described above. Four weeks after PMX, moderate joint damage was accompanied by an increased NaV1.8 signal in the superficial and deep layers of the medial synovium, and no further innervation changes were detected by week 12 *post* PMX (Fig. 2E-H). Representative sections at week 4 are shown in Fig. 2F-H, and synovial innervation at 12-week *post* PMX is shown in Suppl Fig. 6G,H. The innervation of the lateral synovium in the PMX model also declined with age in both sham and PMX groups by week 12, with no evidence of new innervation (Suppl. Fig. 7D).

We selected the ACLR model as a non-invasive injury model of PTOA and assessed joint innervation 1 and 4 weeks after injury. Despite the mild cartilage degeneration at the early 1-week timepoint, fine NaV1.8 fibers had already sprouted into deep and superficial layers of the medial synovium (Fig. 2I-L). No further changes in the medial synovial innervation were detected 4 weeks after injury compared to the early timepoint (Fig. 2I and Suppl. Fig. 6K,L), despite the development of severe cartilage degeneration by this timepoint, especially at the femoral condyles. The innervation in the lateral synovium did not change between week 1 and week 4 in either treatment group (Suppl. Fig. 7G).

Finally, immunostaining for PGP9.5 was used to evaluate neuronal sprouting in knees collected from naïve male C57BL/6 mice at the ages of 6 and 24 months. Six-month old mice showed no nociceptor sprouting in the synovium, concordant with the absence of OA degenerative changes (Suppl. Fig. 2J-L). By 2 years of age, mild joint damage (as observed in Suppl. Fig. 2J) was accompanied by an increase in the sensory innervation of the deep layers of the medial synovium (Fig. 2M-P), but no significant changes in innervation of the lateral synovium were observed between 6 and 24 months (Suppl. Fig. 7J).

#### Nociceptors sprout in subchondral bone channels and within osteophytes

We previously reported the presence of NaV1.8+ nociceptors in channels in the sclerotic subchondral bone, 16 weeks after DMM (16). Here, we analyzed additional timepoints after DMM, and counted the number of medial and lateral channels with a nociceptor signal, as well as the length of the longest channel. This revealed sprouting of NaV1.8+ nerve fibers within medial tibial and femoral subchondral bone channels, 4, 8 and 16 weeks after DMM. We detected no difference in the number or length of channels between the 2 early timepoints (4 and 8 weeks) (Fig. 3A,B). However, the number of NaV1.8 positive channels increased 16 weeks after DMM, as well as the length of the longest channel (Fig. 3A,B) (for the number of channels DMM 4 *vs.* 16 weeks p=0.09, for the length DMM 4 *vs.* 16 weeks p=0.07, one-way ANOVA). Representative images of these channels at 8 and 16 weeks after DMM are shown in Fig. 3C-G. We also assessed the distance from the tidemark and the number of branching points in the nerves within the channels. NaV1.8+ channels were closer to the tidemark and more branched 16 weeks *post* DMM compared to the earlier timepoints (Suppl. Fig. 8A,I) (for the distance from the tidemark DMM 4 *vs.* 16 weeks p=0.027, one-way ANOVA). Representative images at week 4 are shown in Suppl. Fig. 6C,D. Lateral subchondral bone channels were not different between DMM and age-matched shams in any of the parameters measured (Suppl. Fig. 7B,C and Suppl. Fig. 8E). In addition to sprouting in subchondral bone channels, we also detected NaV1.8+ signal within osteophytes at all timepoints after DMM (Fig. 4A-C).

**Figure 3:**
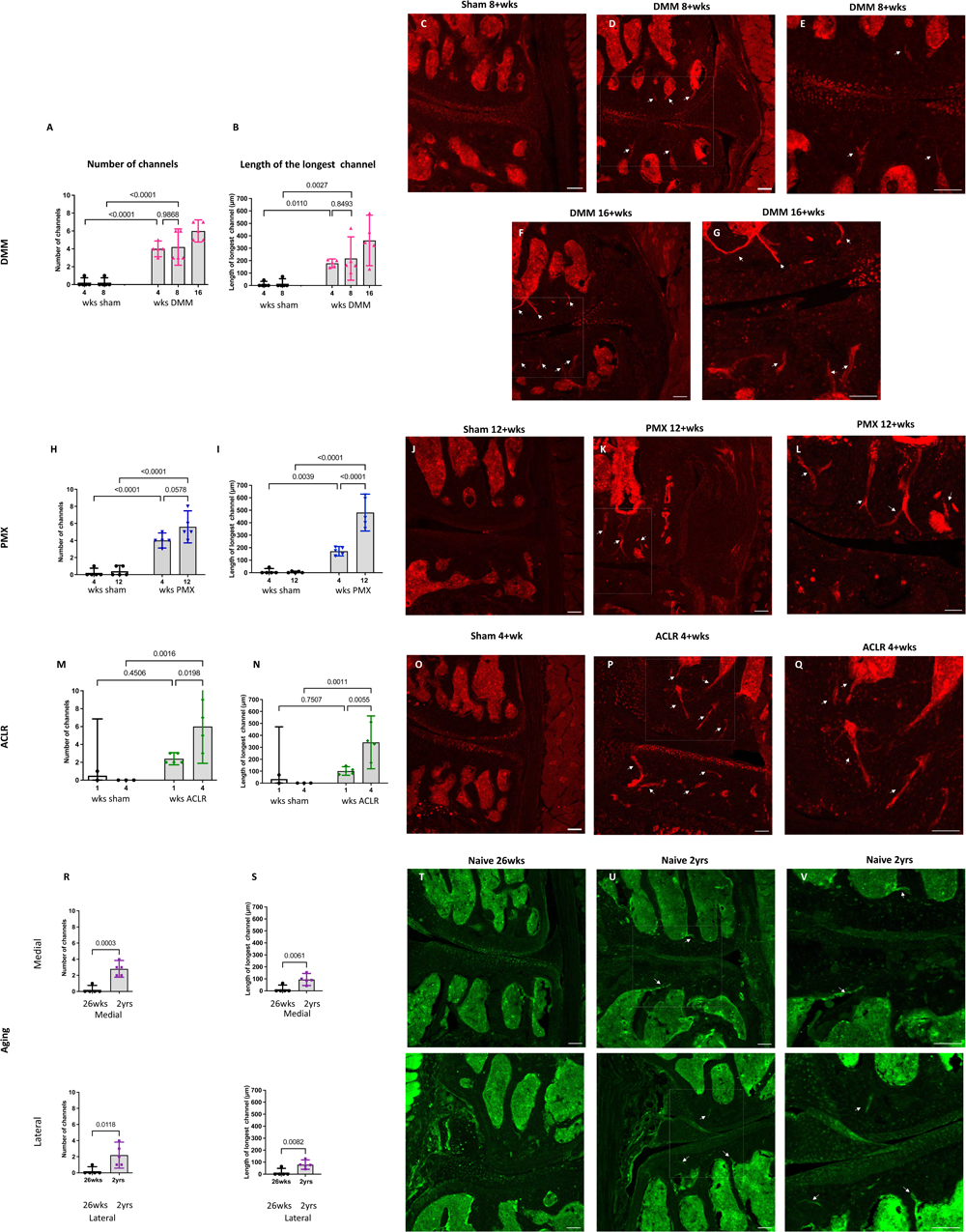
Quantification of neuronal signal in the medial subchondral bone 4, 8 and 16 weeks after sham and DMM surgeries showing (A) the number of NaV1.8+channels; (B) the length of the longest channel; (C-E) Representative confocal images of NaV1.8-tdTomato mouse knees showing fibers within medial subchondral bone channels (white arrows) at 8 weeks after sham and 8 and 16 weeks after DMM surgery; (E,G) magnified image of medial subchondral bone fibers at 8 and 16 weeks after DMM; (H,I) quantification of the number of positive channels and the length of the longest channel respectively at 4 and 12 weeks after sham and PMX surgeries; (J-L) 4 weeks after sham, 4 weeks after PMX surgery, and zoomed in image, respectively; (M,N) quantification of the number of positive channels and the length of the longest channel, respectively; (O-Q) 1 week after sham, 1 week after ACLR injury, and zoomed in image, respectively; (R,S) quantification of the number of positive channels and the length of the longest channel, respectively, in 26-week old and 2-year old naïve mice in the medial and lateral compartment; (T-V) PGP9.5 staining in the medial and lateral compartment of 26-week old mice, 2-year old naïve mice, and zoomed in image, respectively. Mean ± 95% CI. Scale bar = 100 μm.

**Figure 4:**
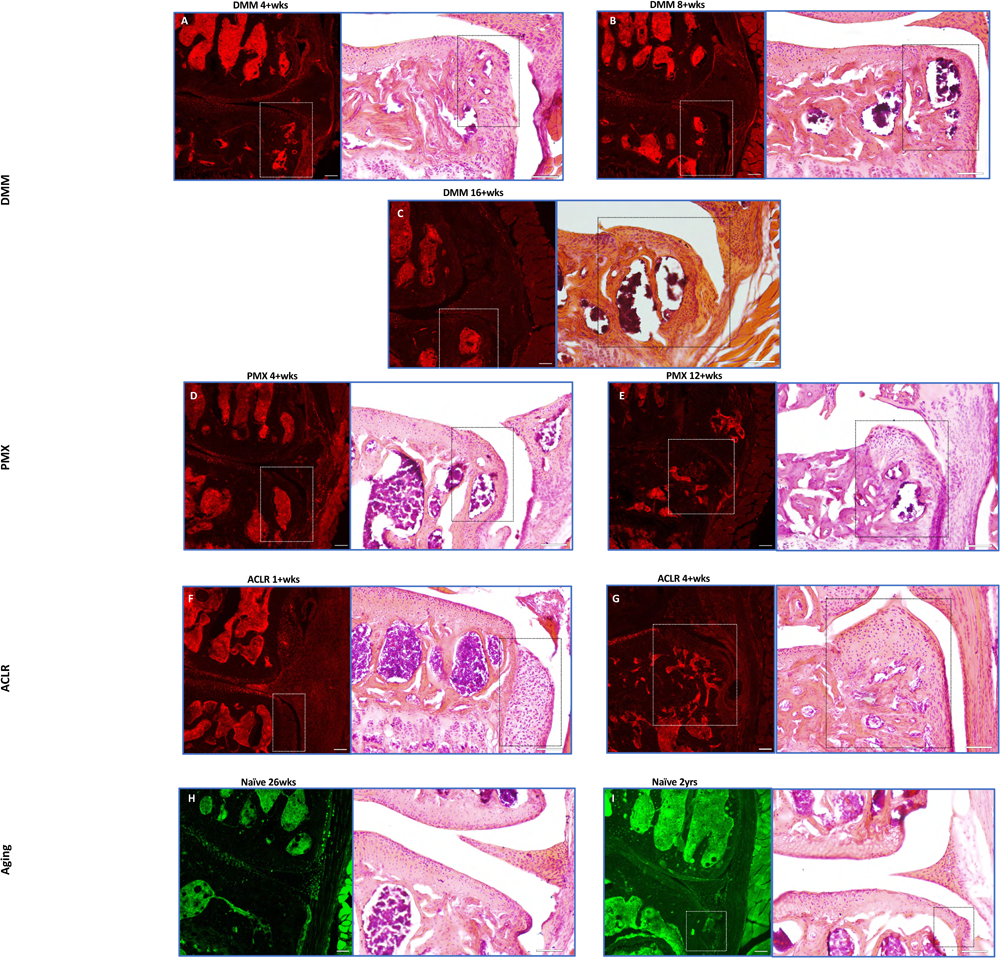
Representative confocal and H&E stained images of NaV1.8-tdTomato and PGP9.5-stained WT knees showing neuronal signal within osteophytes (white and black insets respectively) at (A-C) 4,8 and 16 weeks after DMM; (D,E) 4 and 12 weeks after PMX; (F-G) 1 and 4 weeks after ACLR; (H,I) 26-week old and 2-year old naïve mice. Scale bar = 100 μm.

Similar to DMM, PMX-operated mice showed a significant increase in number of positive channels compared to age-matched sham controls (Fig. 3H). These channels were already prominent by week 4, and by week 12 *post* PMX, NaV1.8+ channels appeared to increase in number (Fig. 3H), were longer (Fig. 3I), closer to the tidemark (Suppl. Fig. 8B), and more branched (Suppl. Fig. 8J), compared to the earlier timepoint. Here too, lateral subchondral bone channels were not different between PMX and age-matched shams in any of the parameters measured (Suppl. Fig. 7E,F and Suppl. Fig. 8F). Representative images showing NaV1.8+ subchondral bone channels at 12+weeks are shown in Fig. 3J-L, and at 4+weeks *post* PMX in Suppl. Fig. 6I,J. We also detected a strong NaV1.8+ signal within osteophytes at both timepoints after PMX (Fig. 4D,E).

One week after ACLR, the subchondral bone showed mild changes, with few Nav1.8+ fibers detectable within channels (Suppl. Fig. 6M,N). By 4 weeks after injury, however, NaV1.8+ subchondral bone channels were observed, and these were significantly greater in number (Fig. 3M), longer (Fig. 3N), and closer to the tidemark (Suppl. Fig. 8C) compared to shams and to the early timepoint. No difference in number and length of lateral channels was observed between ACLR and sham groups (Suppl Fig. 7H,I). Representative images showing NaV1.8+ channels at 4 weeks are shown in Fig. 3O-Q, and at 1 week after ACLR injury in Suppl Fig. 6K,L. No branching was observed at either timepoint (Suppl. Fig. 8K). Mature osteophytes showed NaV1.8+ signal 4 weeks *post* ACLR, but the chondrophytes at the earlier timepoints did not contain nociceptors (Fig. 4F,G).

Finally, PGP9.5 staining of naive knees showed no sprouting in 26-week old mice, when there were no signs yet of primary OA. By age 2 years, OA joint damage in both joint compartments was accompanied by innervation changes, with PGP9.5+ channels present in subchondral bone in both the medial and lateral compartments (Fig. 3T-V). These channels were significantly greater in number, and longer compared to the younger counterparts (Fig. 3R,S). The medial and lateral channels were closer to the tidemark compared to controls (Suppl. Fig. 8D,H). No branching was detected in either compartment. Innervation changes were more pronounced in the medial compartment than in the lateral compartment, reflecting the more pronounced joint changes in the medial compartment. In some old mice, a PGP9.5+ signal was detected within small osteophytes (Fig. 4H,I).

## DISCUSSION

Motivated by increasing evidence that OA of the knee is accompanied by structural plasticity of intra-articular nociceptors, in human subjects as well as in experimental models (15–18), we performed a qualitative and quantitative analysis of the joint nociceptive innervation in 4 mouse models of knee OA. We documented the anatomical distribution of knee joint nociceptors in two surgical models, a non-invasive model of PTOA, and in primary age-associated OA, and this in early and late-stage joint disease.

The key finding in this study is that nociceptor sprouting in the synovium and in subchondral bone channels occurred across all 4 models, in a manner that is tightly linked to joint pathology. In both surgical models and in the non-invasive PTOA model, OA joint damage was restricted to the medial compartment, as was nociceptor sprouting. In contrast, primary age-associated OA affected both compartments, with the medial compartment more severely affected. This was mirrored by nociceptor sprouting, which occurred in both compartments but was more pronounced in the medial compartment.

In the synovial membrane, neo-innervation occurred in a temporal manner that matched the time-course of pathology in the tissue. Nociceptor sprouting had plateaued by the earlier timepoint (week 4 in the surgical models and week 1 after ACLR), consistent with peak synovitis in the early stages of these models. This is consistent with increased nociceptive innervation reported in rodent surgical models (15,16). In contrast, in human OA synovium, a reduction in PGP9.5 and CGRP immunopositive fibers has been described, close to the lining layer (20). In all three induced models, synovitis decreased by the later timepoints, while nociceptor density remained unchanged. Synovitis scores have been previously reported to peak early on in these models, including 4 weeks after DMM (36), as well as in two models of ACL injury, surgically anterior cruciate ligament transection (ACLT) and ACLR (37). Finally, old mice showed mild synovial changes compared to younger mice, confirming our previous findings (25). This was again paralleled by neuronal sprouting in the synovium in these mice.

Neuronal sprouting within subchondral bone channels started at earlier stages, but in contrast to synovial sprouting, this sprouting progressed with worsening joint damage. We detected nerve fibers in subchondral bone channels, and these positive channels were significantly greater in number, longer, and closer to the tidemark at later timepoints compared to earlier timepoints and age-matched sham controls. Changes in subchondral bone innervation were previously reported in a rat medial meniscectomy model, where CGRP-immunopositive nerve fibers within osteochondral channels were detected as early as 2 weeks after surgery, and increased in density at later stages (15). Subchondral bone neuroplasticity was first reported in human OA knees, where CGRP immunopositive fibers were detected within osteochondral channels, and – as in the rat-were associated with pain (13). Interestingly, we found that severe osteoarthritic changes in late-stage OA were associated with more branching of sprouted nerves within bone channels, which may result in an increased receptive field, as has been described in the skin (38).

Finally, all 4 models showed abundant nociceptors within bone marrow cavities in osteophytes. Sprouting was only observed in relatively mature, mineralized osteophyte, not in the precursor chondrophytes. This indicates that mineralization, angiogenesis, and nerve sprouting are potentially coupled processes in the context of osteophytes. PGP9.5 nerve trunks have also been described in subchondral bone marrow and within osteophyte marrow cavities in knees after joint replacement surgery for tibiofemoral OA (39).

In short, our findings suggest that experimental OA is accompanied by profound neuronal remodeling in different joint tissues, starting early after OA induction and associated with disease progression. In the three induced models, joint damage occurred predominantly in the medial compartment of the knee, as expected, and neuronal sprouting was also confined to this compartment. Of note, in primary OA in ageing mice, joint damage occurred in both the medial and lateral compartment, and this was accompanied by neuronal sprouting in both compartments although it was greater in the medial compartment, reflecting the more severe joint damage in that compartment. These observations, together with reports on nociceptor sprouting in human knees, suggest that nociceptor sprouting is a consistent anatomical hallmark of OA joint damage, and that it is intrinsic to the disease. It also appears that neuronal sprouting starts early in the disease, both in the synovium and in the subchondral bone, and is established more rapidly in the synovium than in the bone. This is concordant with a recent study in rats, which reported that CGRP+ synovial innervation was increased at early stages (2 weeks after medial meniscectomy) and then decreased at later stages, while the sprouting in the subchondral bone continued to increase over time (15). In older mice, where we have previously reported that the existing nociceptor innervation in the lateral synovium declines between the ages of 10 weeks and of 26 weeks (16), we observed that by age 2 years, sprouting accompanied mild OA joint damage.

These observations raise the question as to what the biological role of this neuronal sprouting might be, and how it relates to joint damage and pain. Hence, it would be reasonable to conclude that NaV1.8+ neo-innervation contributes to pain hypersensitivity in the knee, but demonstrating this will require sophisticated approaches to selectively ablate or silence these particular neurons, and assess the effect on pain. Our findings might be discussed in the context of a study that utilized *in vivo* electrophysiology to document responses of knee-innervating and bone-innervating neurons in the rat monosodium iodoacetate (MIA) model, which compared responses at an early (day 3) and a late timepoint (day 28) (40). This study reported that early pain involved activation and sensitization of nerves within the medial aspect of the joint capsule, which is lined with the synovium where we detected substantial neuronal sprouting at early stages of the disease across all models. Moreover, the authors reported that pain in the late-stage MIA model was associated with recruitment of nerves in the subchondral bone (40). In addition, a study in knees of patients undergoing total knee replacement for painful knee OA reported increased CGRP+ osteochondral channels compared to asymptomatic *post-*mortem controls, suggesting that these nociceptors in the subchondral bone contribute to pain in late-stage OA (13).

Secondly, sensory joint innervation might also be important for joint homeostasis and maintenance of joint health. Loss of joint afferents (including large myelinated nerves) leads to marked destruction of articular joints in a condition called neuropathic joints or Charcot’s arthropathy, such as in diabetic neuropathy (41). There is also evidence from animal models that crude denervation may cause joint damage. For example, joint denervation through unilateral dorsal root ganglionectomy in the canine ACLT model led to accelerated cartilage degeneration (42). A similar study used retrogradely transported immunotoxin to denervate rat knees, and this resulted in severe osteoarthritic changes (43). This evidence from human disease and animal models suggests that intra-articular sensory neurons play a critical role in joint health, as opposed to just mediating nociceptive and proprioceptive responses. More work is needed to determine the specific role of nociceptors and nociceptor sprouting, an evidently early phenomenon in experimental OA, but we speculate that their role extends beyond pain, and they may well have an efferent role that is critical for joint homeostasis after injury. For example, free nerve endings release inflammatory neuropeptides such as CGRP and substance P, which cause vasodilation and leukocyte extravasation (44), and these might enable multiple other tissue changes within the joint. In order to explore the biological role of joint nociceptor plasticity, it will be critical to identify the factors that drive it. One potential candidate to explain nerve sprouting in OA models is the neurotrophin, NGF, which has been shown to promote nerve sprouting in bone cancer and inflammatory models (45,46). NGF is expressed in OA joints, in the synovium and in osteochondral channels (15,47,48). Clinical trials with tanezumab, a monoclonal antibody that neutralizes NGF, showed promising results in treating OA pain but the occurrence of rapidly progressive OA in some patients halted the development of these antibodies. The role of NGF in joint nociceptor sprouting and in joint homeostasis needs to be explored. Our early findings suggest that intra-articular administration of NGF cause synovitis and nociceptor sprouting in the medial synovium and subchondral bone (49). Future studies will explore the functional significance of newly sprouted nerves by selective ablation of knee nociceptors and study the effect on joint integrity and OA pain.

A notable limitation of our study is the fact that all experiments were only conducted in male mice. Ongoing studies in the PMX and in the ACLR model suggest that sprouting also occurs in female mice (26), and future studies will investigate sex-specific processes in these models.

In summary, nociceptor sprouting was observed in specific knee joint tissues in 4 experimental models of OA, occurring in a specific temporal manner in association with progressive joint pathology. The tight relationship between joint pathology and neuronal sprouting suggests that neuroplasticity is a fundamental component of OA joint pathology. Together with the reports of neuronal plasticity in human knee joints, our findings warrant an in-depth exploration of the innervation of the OA knee joint, the drivers of the observed neuroplasticity, and the communication between joint nociceptors and the joint tissues they innervate. It can be expected that a precise elucidation of these processes will lead to new targets for OA pain as well as joint homeostasis.

## Supporting information

Suppl. Methods, Suppl. Results

## Acknowledgements

The authors would like to thank Dr. Matthew Wood for his help in making Figure 1.

## Supplemental Figure legends

**Suppl. Figure 1:**
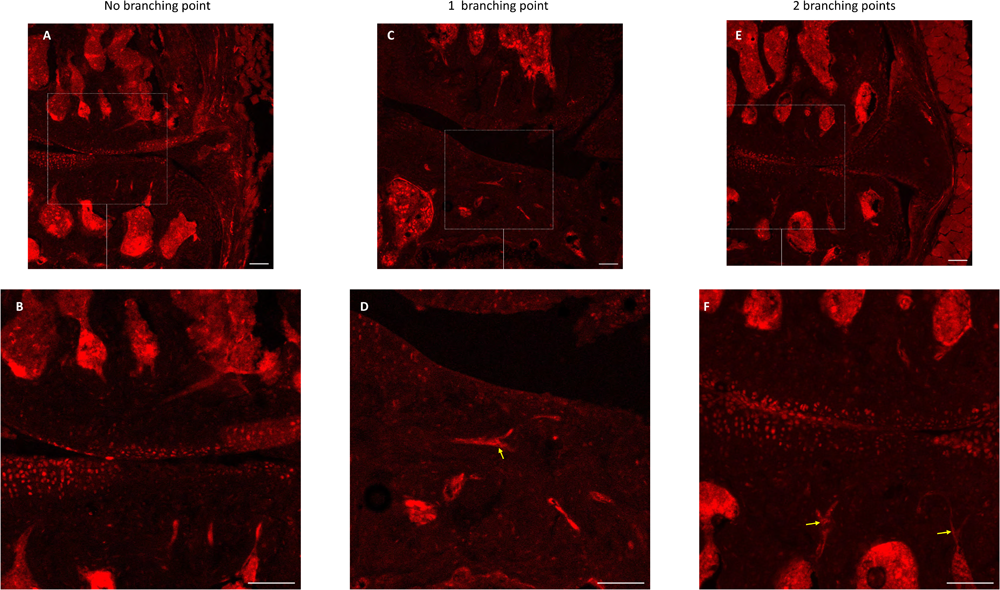
Quantification of branching points in NaV1.8+ subchondral bone channels. A and B (zoomed in) show confocal images with no branching points; C and D show an image with one branching point, indicated by the yellow arrow; E and F show a confocal image with two branching points, indicated by the yellow arrows. Scale bare =100 μm.

**Suppl. Figure 2:**
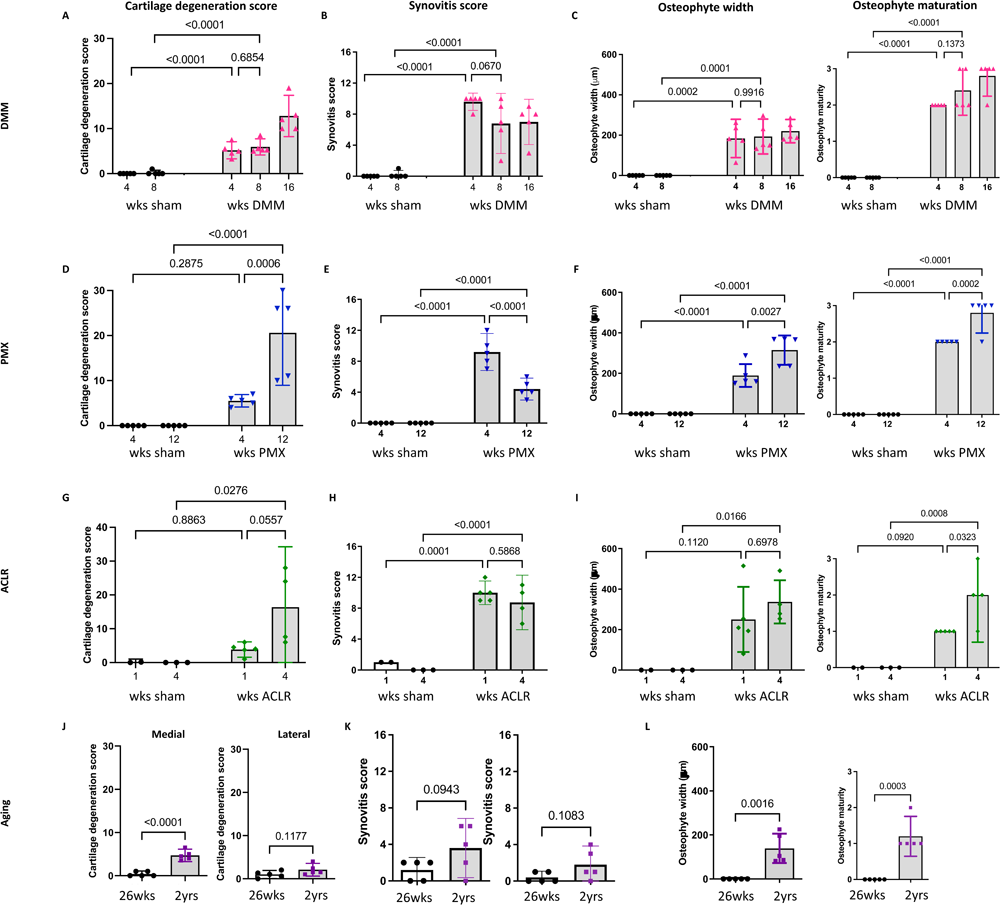
(A-C) Medial cartilage degeneration scores, synovitis scores, osteophyte width (measured in μm), and osteophyte maturity in right knees of 4, 8 and 16 weeks after sham or DMM surgery in male mice; (D-F) Scores in right knees 4 and 12 weeks after sham or PMX in male mice; (G-I) Scores in right knees 1 and 4 weeks after sham or ACLR in male mice; (J-L) Scores in right knees of 26-week old and 2-year old naïve male mice. Wks=weeks. Medial cartilage degeneration DMM 4+wks = 5.18±1.5, sham 4+wks = 0; DMM 8+wks = 6±1.4, sham 8+wks = 0.2±0.4; DMM 16+wks = 12.8±3.7; PMX 4+wks = 5.6±1.2, sham 4+wks = 0, PMX 12+wks = 20.6±9.3, sham 12+wks = 0; ACLR 1+wk = 3.85±1.8, sham 1+wk = 0.07±0.1; ACLR 4+wks = 16.4±11.2, sham 4+wks = 0. Medial cartilage degeneration score for 26-wk old mice = 0.4±0.55, and lateral cartilage degeneration score for 26-wk old mice = 0.98±0.74; Medial cartilage degeneration score for 2-year old mice = 4.72±1.19, and lateral = 2.08±1.18. Synovitis scores are sum scores of synovial hyperplasia, cellularity, and fibrosis for the medial compartment of the right knees. Osteophyte width was measured in μm for the major osteophyte in the medial compartment. Osteophyte width for DMM 4 weeks= 183.9±76.6, DMM 8 weeks= 193.2±69.6, PMX 4 weeks= 189.2±56.6, PMX 12 weeks= 314.7±72.7, ACLR 1 week= 250.2±161.3, ACLR 4 weeks= 336.9.±106.5, 2-year old naïve mice= 138.8±66.8. All graphs show the mean ± 95% CI.

**Suppl. Figure 3:**
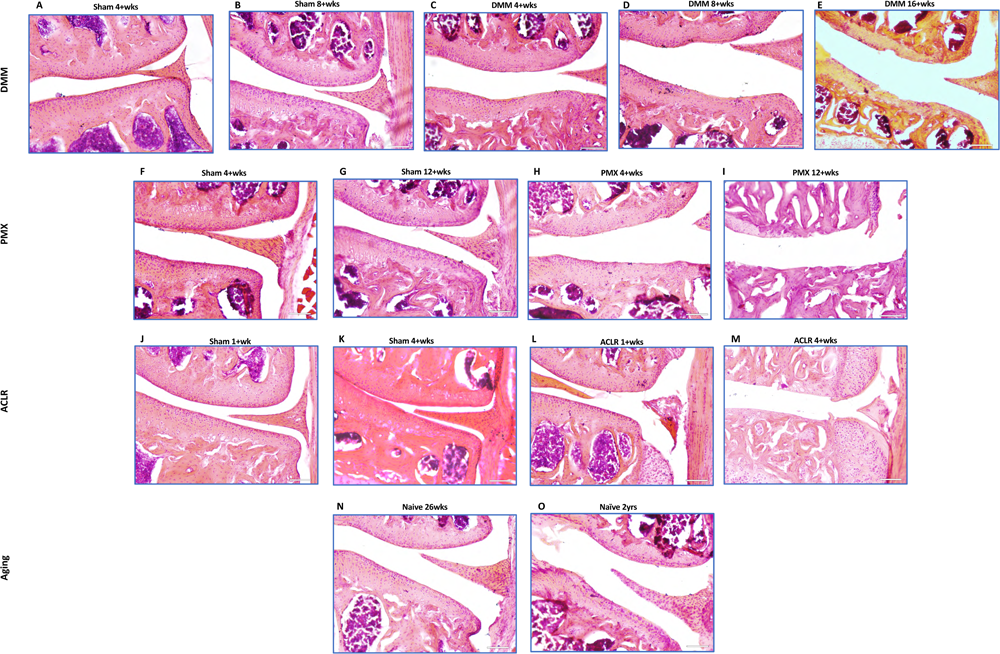
Representative histological images of the medial compartment of NaV1.8-tdTomato and WT knees (A-E) at 4 and 8 after sham and 4, 8 and 16 after DMM surgery; (F-I) 4 and 12 weeks after sham or PMX surgery; (J-M) 1 and 4 weeks after sham or ACLR; (N,O) knees from 26 week-old and 2 year-old naïve mice. Scale bar = 100 μm.

**Suppl. Figure 4:**
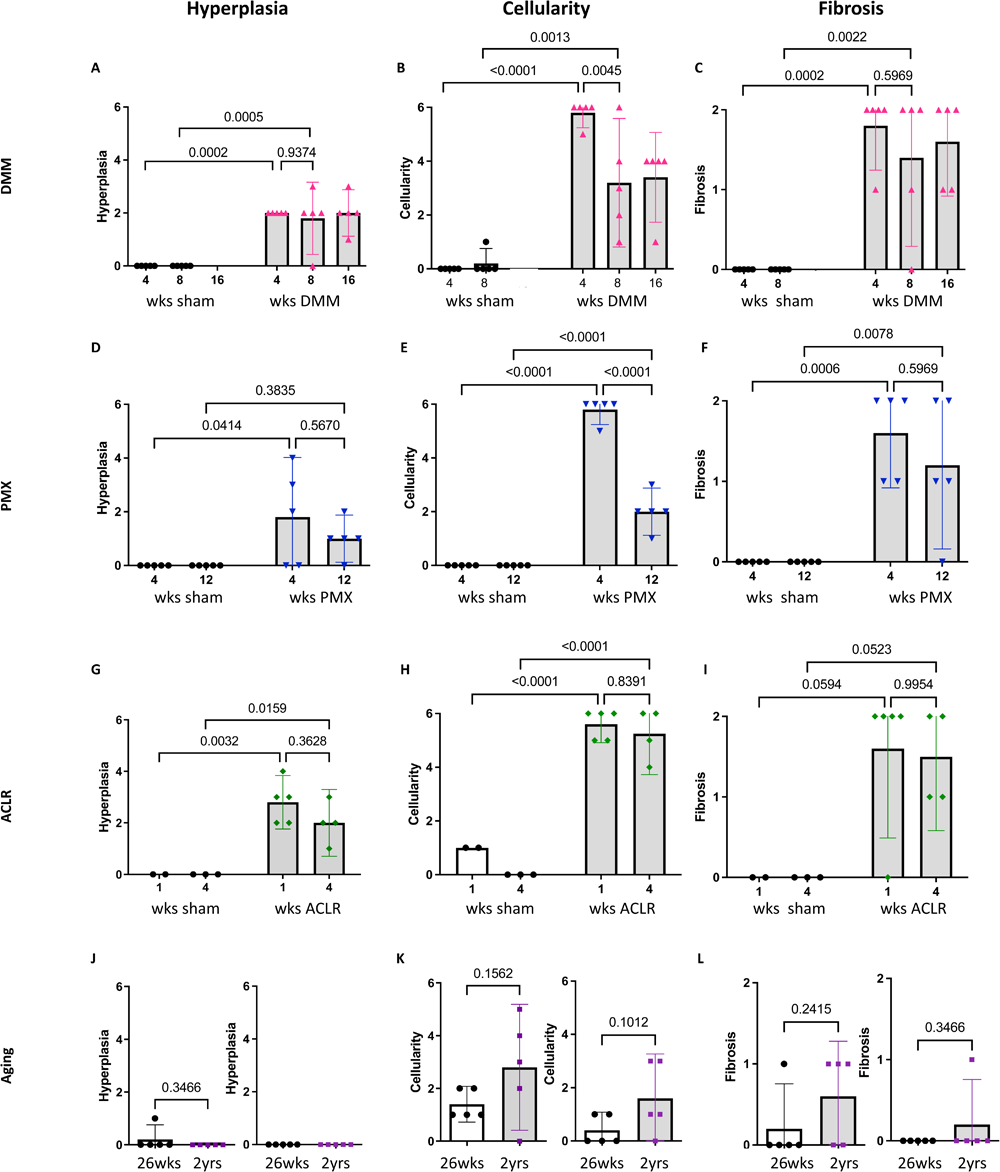
Synovial scoring of hyperplasia, cellularity and fibrosis, respectively, including total scores from medial femoral and medial tibial joint spaces. (A-C) 4, 8 and 16 weeks after sham or DMM surgery; (D-F) 4 and 12 weeks after sham or PMX surgery; (G-I) 1 and 4 weeks after sham or ACLR injury; (J-L) 26-week old and 2-year old naïve mice. Mean ± 95% CI.

**Suppl. Figure 5:**
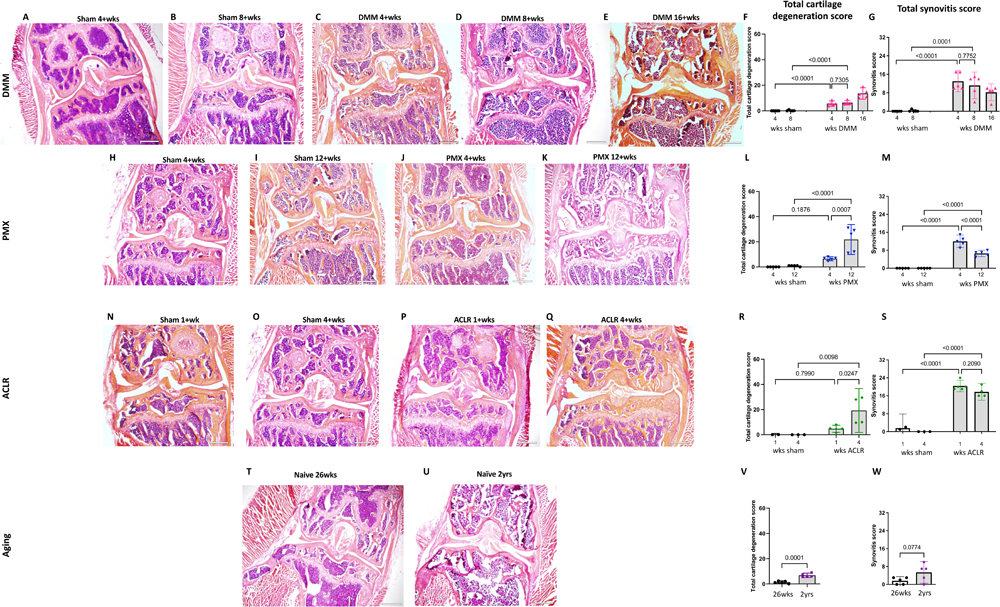
Representative histological images of whole knees of NaV1.8-tdTomato and WT mice (A-E) 4, 8, and 16 weeks after sham or DMM surgery; (F,G) total cartilage degeneration score and synovitis total limb score, respectively; (H-K) 4 and 12 weeks after sham or PMX surgery; (L,M) total cartilage degeneration score and synovitis total limb score, respectively; (N-Q) 1 and 4 weeks after sham or ACLR injury; (R,S) total cartilage degeneration score and synovitis total limb score, respectively; (T-U) knees from 26-week old and 2-year old naïve mice; (V,W) total cartilage degeneration score and synovitis total limb score, respectively. Mean ± 95% CI. Scale bar = 500 μm.

**Suppl. Figure 6:**
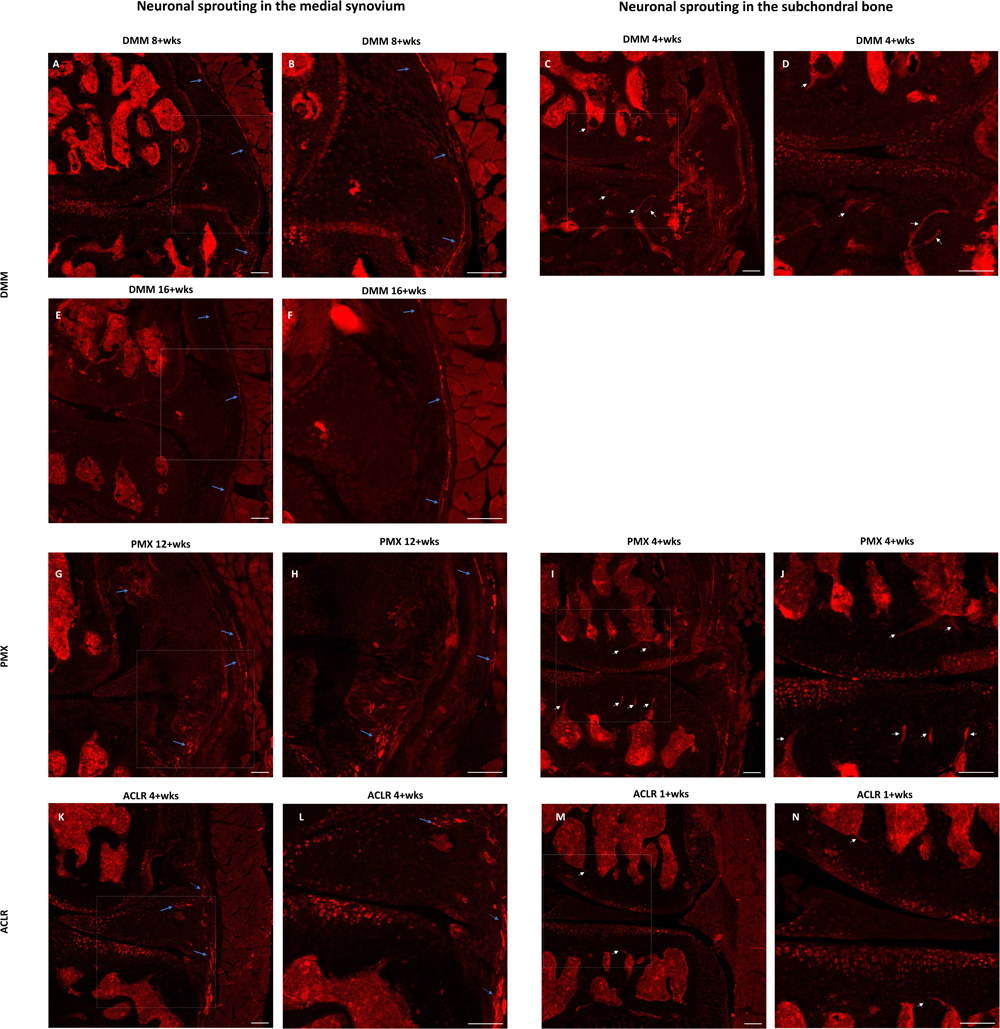
Representative confocal images of NaV1.8-tdTomato mouse knees showing (A,B,E,F) neoinnervation in the medial synovium (blue arrows) 8 and 16 weeks after sham and DMM surgery; (C,D) nerve fibers in subchondral bone channels (white arrows) 4 weeks after sham and DMM surgery; (G,H) nerve fibers in the medial synovium 12 weeks after sham and PMX surgery; (I,J) nerve fibers within subchondral bone channels 4 weeks after sham and PMX surgery; (E,L) nerve fibers in the medial synovium 4 weeks after sham and ACLR injury; (M,N) nerve fibers within subchondral bone channels 1 week after sham and ACLR injury. Scale bar = 100 μm.

**Suppl. Figure 7:**
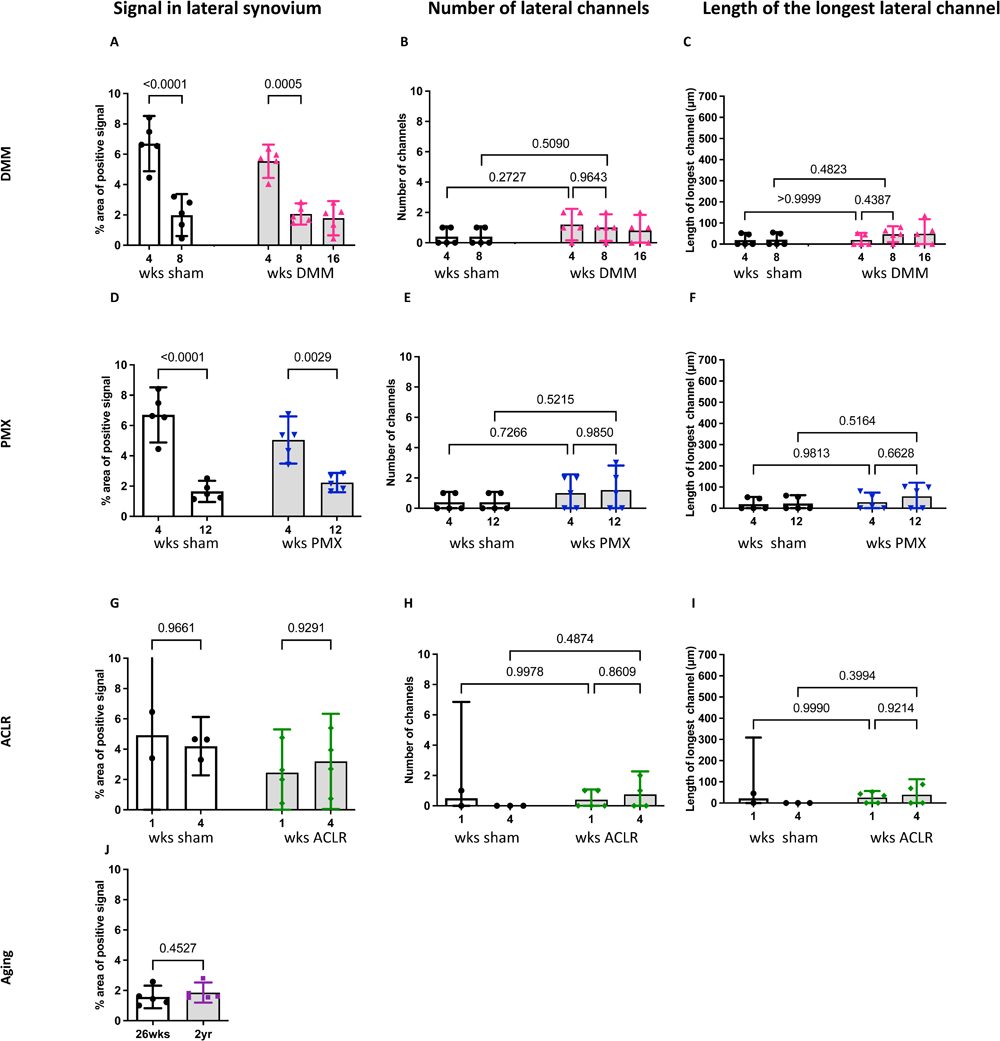
Quantification of NaV1.8+ signal in the lateral synovium, the number of lateral subchondral bone channels and the length of the longest channel at (A-C) 4, 8 and 16 weeks after DMM or sham surgery; (D-F) 4 and 12 weeks after PMX or sham surgery; (G-I) 1 and 4 weeks after ACLR injury or sham; (J) 26-week old and 2-year old naïve mice. Mean ± 95% CI.

**Suppl. Figure 8:**
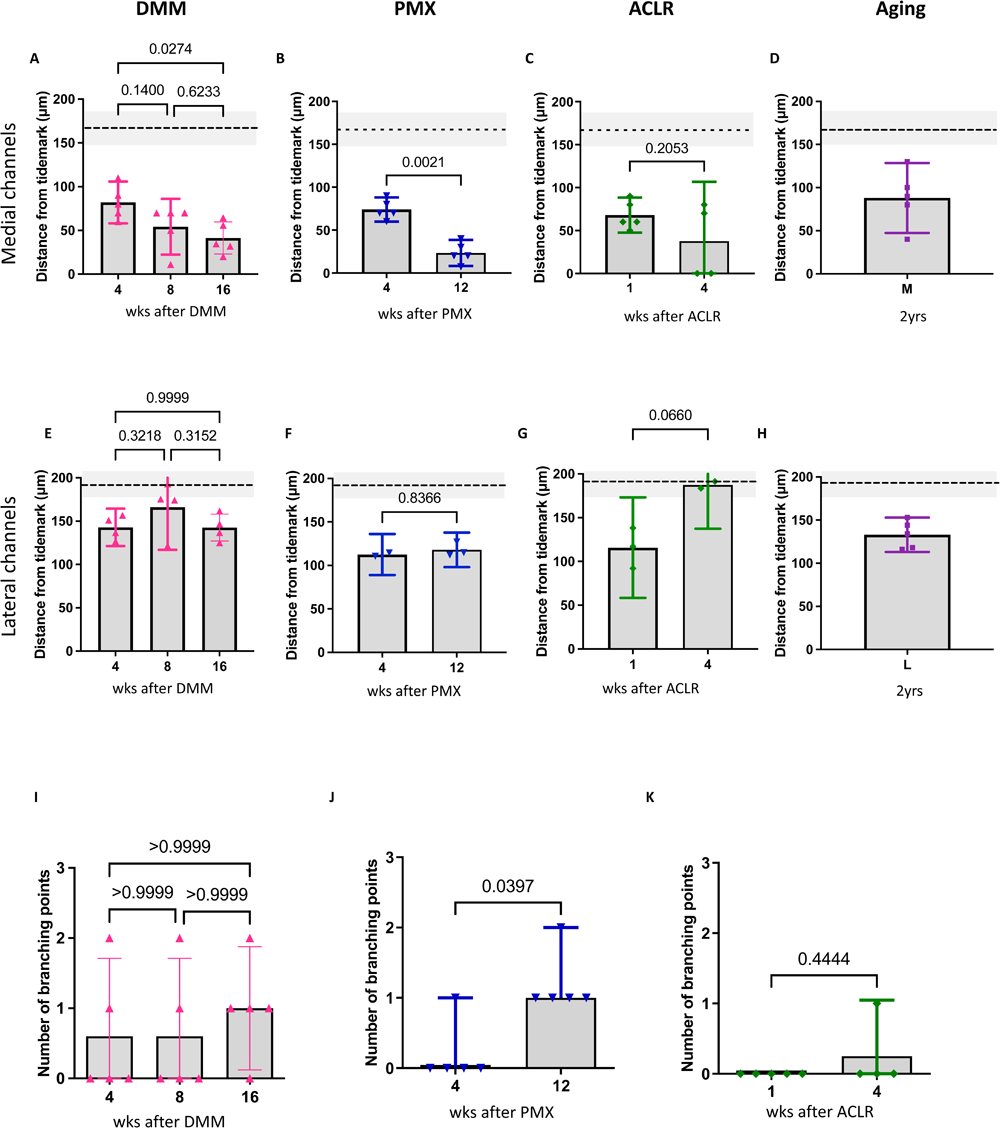
(A-D) Quantification of the distance of NaV1.8+ and PGP9.5+ medial channels from the tidemark; (E-H) Quantification of the distance of NaV1.8+ and PGP9.5+ lateral channels from the tidemark; (I-K) Number of branching points 4, 8 and 16 weeks after DMM, 4 and 12 weeks after PMX surgery, and 1 and 4 weeks after ACLR injury. M=medial, L=lateral; (A-H) Mean ± 95% CI. (I-K) Median ± 95% CI.

## REFERENCES

1. Kraus VB, Blanco FJ, Englund M, Karsdal MA, Lohmander LS. Call for standardized definitions of osteoarthritis and risk stratification for clinical trials and clinical use. Osteoarthritis Cartilage. 2015 Aug;23(8):1233–41.

2. Loeser RF, Goldring SR, Scanzello CR, Goldring MB. Osteoarthritis: a disease of the joint as an organ. Arthritis Rheum. 2012 Jun;64(6):1697–707.

3. Latourte A, Kloppenburg M, Richette P. Emerging pharmaceutical therapies for osteoarthritis. Nat Rev Rheumatol. 2020 Dec;16(12):673–88.

4. Bedson J, Mottram S, Thomas E, Peat G. Knee pain and osteoarthritis in the general population: what influences patients to consult? Fam Pract. 2007 Oct;24(5):443–53.

5. Neogi T. The epidemiology and impact of pain in osteoarthritis. Osteoarthritis Cartilage. 2013 Sep;21(9):1145–53.

6. Deveza LA, Hunter DJ, Van Spil WE. Too much opioid, too much harm. Osteoarthritis Cartilage. 2018 Mar;26(3):293–5.

7. Vincent TL. Peripheral pain mechanisms in osteoarthritis. Pain. 2020 Sep;161 Suppl 1(1):S138–46.

8. Malfait AM, Miller RE, Miller RJ. Basic Mechanisms of Pain in Osteoarthritis. Rheum Dis Clin N Am. 2021 May;47(2):165–80.

9. Wise BL, Seidel MF, Lane NE. The evolution of nerve growth factor inhibition in clinical medicine. Nat Rev Rheumatol. 2021 Jan;17(1):34–46.

10. Schaible HG, Grubb BD. Afferent and spinal mechanisms of joint pain. Pain. 1993 Oct;55(1):5–54.

11. Heppelmann B. Anatomy and histology of joint innervation. J Peripher Nerv Syst JPNS. 1997;2(1):5– 16.

12. McDougall JJ. Arthritis and pain. Neurogenic origin of joint pain. Arthritis Res Ther. 2006;8(6):220.

13. Aso K, Shahtaheri SM, Hill R, Wilson D, McWilliams DF, Nwosu LN, et al. Contribution of nerves within osteochondral channels to osteoarthritis knee pain in humans and rats. Osteoarthritis Cartilage. 2020 Sep;28(9):1245–54.

14. Ishihara S, Obeidat AM, Wokosin DL, Ren D, Miller RJ, Malfait AM, et al. The role of intra-articular neuronal CCR2 receptors in knee joint pain associated with experimental osteoarthritis in mice. Arthritis Res Ther. 2021 Apr 7;23(1):103.

15. Aso K, Walsh DA, Wada H, Izumi M, Tomitori H, Fujii K, et al. Time course and localization of nerve growth factor expression and sensory nerve growth during progression of knee osteoarthritis in rats. Osteoarthritis Cartilage. 2022 Oct;30(10):1344–55.

16. Obeidat AM, Miller RE, Miller RJ, Malfait AM. The nociceptive innervation of the normal and osteoarthritic mouse knee. Osteoarthritis Cartilage. 2019 Nov;27(11):1669–79.

17. Buma P, Verschuren C, Versleyen D, Van der Kraan P, Oestreicher AB. Calcitonin gene-related peptide, substance P and GAP-43/B-50 immunoreactivity in the normal and arthrotic knee joint of the mouse. Histochemistry. 1992 Dec;98(5):327–39.

18. Murakami K, Nakagawa H, Nishimura K, Matsuo S. Changes in peptidergic fiber density in the synovium of mice with collagenase-induced acute arthritis. Can J Physiol Pharmacol. 2015 Jun;93(6):435–41.

19. Ter Heegde Fs, Luiz AP, Santana-Varela S, Magnúsdóttir R, Hopkinson M, Chang Y, et al. Osteoarthritis-related nociceptive behaviour following mechanical joint loading correlates with cartilage damage. Osteoarthritis Cartilage. 2020 Mar;28(3):383–95.

20. Eitner A, Pester J, Nietzsche S, Hofmann GO, Schaible HG. The innervation of synovium of human osteoarthritic joints in comparison with normal rat and sheep synovium. Osteoarthritis Cartilage. 2013 Sep;21(9):1383–91.

21. Aso K, Shahtaheri SM, Hill R, Wilson D, McWilliams DF, Walsh DA. Associations of Symptomatic Knee Osteoarthritis With Histopathologic Features in Subchondral Bone. Arthritis Rheumatol. 2019 Jun;71(6):916–24.

22. Knights CB, Gentry C, Bevan S. Partial medial meniscectomy produces osteoarthritis pain-related behaviour in female C57BL/6 mice. Pain. 2012 Feb;153(2):281–92.

23. Rzeczycki P, Rasner C, Lammlin L, Junginger L, Goldman S, Bergman R, et al. Cannabinoid receptor type 2 is upregulated in synovium following joint injury and mediates anti-inflammatory effects in synovial fibroblasts and macrophages. Osteoarthritis Cartilage. 2021 Dec;29(12):1720–31.

24. Loeser RF. Aging processes and the development of osteoarthritis. Curr Opin Rheumatol. 2013 Jan;25(1):108–13.

25. Geraghty T, Obeidat AM, Ishihara S, Wood MJ, Li J, Lopes EBP, et al. Age-associated changes in knee osteoarthritis, pain-related behaviors, and dorsal root ganglia immunophenotyping of male and female mice. Arthritis Rheumatol. 2023 Apr 25;art.42530.

26. Bergman RF, Lammlin L, Junginger L, Farrell E, Goldman S, Darcy R, et al. Sexual dimorphism of the synovial transcriptome underpins greater PTOA disease severity in male mice following joint injury [Internet]. Physiology; 2022 Dec [cited 2023 Jan 26]. Available from: http://biorxiv.org/lookup/doi/10.1101/2022.11.30.517736

27. von Loga IS, Batchelor V, Driscoll C, Burleigh A, Chia SLL, Stott B, et al. Does Pain at an Earlier Stage of Chondropathy Protect Female Mice Against Structural Progression After Surgically Induced Osteoarthritis? Arthritis Rheumatol Hoboken NJ. 2020 Dec;72(12):2083–93.

28. Miller RE, Malfait AM. Osteoarthritis pain: What are we learning from animal models? Best Pract Res Clin Rheumatol. 2017 Oct;31(5):676–87.

29. Stirling CL, Forlani G, Baker MD, Wood JN, Matthews EA, Dickenson AH, et al. Nociceptor-specific gene deletion using heterozygous NaV1.8-Cre recombinase mice. Pain. 2005 Jan;113(1):27–36.

30. Shields SD, Ahn HS, Yang Y, Han C, Seal RP, Wood JN, et al. Nav1.8 expression is not restricted to nociceptors in mouse peripheral nervous system. Pain. 2012 Oct;153(10):2017–30.

31. Glasson SS, Blanchet TJ, Morris EA. The surgical destabilization of the medial meniscus (DMM) model of osteoarthritis in the 129/SvEv mouse. Osteoarthritis Cartilage. 2007 Sep;15(9):1061–9.

32. Arshadi C, Günther U, Eddison M, Harrington KIS, Ferreira TA. SNT: a unifying toolbox for quantification of neuronal anatomy. Nat Methods. 2021 Apr;18(4):374–7.

33. Little CB, Barai A, Burkhardt D, Smith SM, Fosang AJ, Werb Z, et al. Matrix metalloproteinase 13-deficient mice are resistant to osteoarthritic cartilage erosion but not chondrocyte hypertrophy or osteophyte development. Arthritis Rheum. 2009 Dec;60(12):3723–33.

34. Sekiya I, Sasaki S, Miura Y, Aoki H, Katano H, Okanouchi N, et al. Medial Tibial Osteophyte Width Strongly Reflects Medial Meniscus Extrusion Distance and Medial Joint Space Width Moderately Reflects Cartilage Thickness in Knee Radiographs. J Magn Reson Imaging JMRI. 2022 Sep;56(3):824– 34.

35. Gilbert SJ, Bonnet CS, Stadnik P, Duance VC, Mason DJ, Blain EJ. Inflammatory and degenerative phases resulting from anterior cruciate rupture in a non-invasive murine model of post-traumatic osteoarthritis: INFLAMMATORY AND DEGENERATIVE CHANGES IN POST-TRAUMATIC OSTEOARTHRITIS. J Orthop Res. 2018 Aug;36(8):2118–27.

36. Shu CC, Zaki S, Ravi V, Schiavinato A, Smith MM, Little CB. The relationship between synovial inflammation, structural pathology, and pain in post-traumatic osteoarthritis: differential effect of stem cell and hyaluronan treatment. Arthritis Res Ther. 2020 Feb 14;22(1):29.

37. Urban H, Blaker C, Shu C, Clarke E, Little CB. Synovial inflammation in anterior cruciate ligament injury knees in mice: surgical vs non-surgical models. Osteoarthritis Cartilage. 2020 Apr;28:S213–4.

38. Peng YB, Ringkamp M, Campbell JN, Meyer RA. Electrophysiological assessment of the cutaneous arborization of Adelta-fiber nociceptors. J Neurophysiol. 1999 Sep;82(3):1164–77.

39. Suri S, Gill SE, Massena de Camin S, Wilson D, McWilliams DF, Walsh DA. Neurovascular invasion at the osteochondral junction and in osteophytes in osteoarthritis. Ann Rheum Dis. 2007 Nov;66(11):1423–8.

40. Morgan M, Thai J, Nazemian V, Song R, Ivanusic JJ. Changes to the activity and sensitivity of nerves innervating subchondral bone contribute to pain in late-stage osteoarthritis. Pain. 2022 Feb 1;163(2):390–402.

41. Potts WJ. THE PATHOLOGY OF CHARCOT JOINTS. Ann Surg. 1927 Oct;86(4):596–606.

42. O’Connor BL, Palmoski MJ, Brandt KD. Neurogenic acceleration of degenerative joint lesions. J Bone Joint Surg Am. 1985 Apr;67(4):562–72.

43. Salo PT, Hogervorst T, Seerattan RA, Rucker D, Bray RC. Selective joint denervation promotes knee osteoarthritis in the aging rat. J Orthop Res Off Publ Orthop Res Soc. 2002 Nov;20(6):1256–64.

44. McDougall JJ. Osteoarthritis is a neurological disease – an hypothesis. Osteoarthr Cartil Open. 2019 Nov;1(1–2):100005.

45. Jimenez-Andrade JM, Bloom AP, Stake JI, Mantyh WG, Taylor RN, Freeman KT, et al. Pathological Sprouting of Adult Nociceptors in Chronic Prostate Cancer-Induced Bone Pain. J Neurosci. 2010 Nov 3;30(44):14649–56.

46. Ghilardi JR, Freeman KT, Jimenez-Andrade JM, Coughlin KA, Kaczmarska MJ, Castaneda-Corral G, et al. Neuroplasticity of sensory and sympathetic nerve fibers in a mouse model of a painful arthritic joint. Arthritis Rheum. 2012 Jul;64(7):2223–32.

47. Mapp PI, Avery PS, McWilliams DF, Bowyer J, Day C, Moores S, et al. Angiogenesis in two animal models of osteoarthritis. Osteoarthritis Cartilage. 2008 Jan;16(1):61–9.

48. Stoppiello LA, Mapp PI, Wilson D, Hill R, Scammell BE, Walsh DA. Structural associations of symptomatic knee osteoarthritis. Arthritis Rheumatol Hoboken NJ. 2014 Nov;66(11):3018–27.

49. Obeidat A, Ishihara S, Wang L, Miller R, Malfait A. Effect Of Repeated Injections Of Intra-Articular Nerve Growth Factor In The Naïve Murine Knee Joint. [abstract]. ORS. 2022.

