## Supplementary material for "Intra-Articular Sprouting Of Nociceptors Accompanies Progressive Osteoarthritis: Comparative Evidence In Four Murine Models": Suppl. Methods, Suppl. Results: Supplementary Material .pdf

**Supplementary Methods**

**Destabilization of the medial meniscus (DMM)** – Mice were anesthetized by inhalation of isoflurane. The joint capsule was opened and the anterior medial meniscotibial ligament was severed. The knee was flushed with saline, and the incision closed. Sham surgery was identical to DMM, except that the ligament was left intact (1).

**Partial meniscectomy (PMX)** – Mice were anesthetized by inhalation of isoflurane. After medial parapatellar arthrotomy, the infra-patellar fat pad was dissected to expose the anterior part of the medial compartment of the knee. The medial meniscotibial ligament was transected to release the anterior horn of the medial meniscus, and approximately 1/3 to 1/2 of the anterior portion of medial meniscus was cut. For sham surgery, the medial meniscotibial ligament and the medial meniscus were left intact (2).

**Anterior cruciate ligament rupture (ACLR)** – ACLR mice were subjected to a tibial compression-based noninvasive ACL rupture protocol, as described (3). Briefly, mice were anesthetized with 2% inhaled isoflurane and immobilized on a custom fixture on a mechanical testing system (Electroforce 3300AT, TA Instruments, New Castle, DE). The right knee was flexed to 100$^{o}$ and secured within a trough to restrict medial-lateral motion. The hindpaw was secured in 30$^{o}$ dorsiflexion. After preloading and conditioning, a rapid displacement of 1.5 mm (10 mm/s) was applied via the hindpaw, causing tibial compression, anterior tibial subluxation, and ACL rupture. Complete ACL rupture was confirmed via an anterior drawer test. Following the rupture or sham procedure, animals were administered a single dose of subcutaneous carprofen (5 mg/kg). Mice were allowed *ad libitum* cage activity and provided unlimited access to food and water. Mice were housed in ventilated cages containing a maximum of 5 animals (randomized mix of Sham and ACLR) and maintained within a 12- hour light/dark cycle facility (4).

**Knee histopathology –** Knee sections were evaluated for cartilage degeneration using a modified OARSI score, as described (5). Four joint surfaces, medial and lateral femoral condyles and tibial plateaux were scored for severity of cartilage degeneration. For each cartilage surface, scores were assigned individually to each of three zones (inner, middle, outer) on a scale of 0–5, with 5 representing the most damage. The maximum score for the sum of 60 femoral and tibial cartilage degeneration on either the medial or lateral side = 30. The maximum possible total cartilage degeneration score for the whole joint (sum of medial and lateral sides) is 60.

### Supplementary Results

**Knee histopathology** –We assessed OARSI score, synovitis, and osteophyte width and maturity at different stages of disease.

*OARSI score:* Four and eight weeks after DMM, knees showed mild to moderate cartilage damage (medial cartilage degeneration score for DMM 4+wks = 5.18±1.5, sham 4+wks = 0, DMM 8+wks = 6±1.4, sham 8+wks = 0.2±0.4) (medial cartilage degeneration Suppl. Fig. 2A, and Suppl. Fig. 5F for total cartilage degeneration), Sixteen weeks after DMM, cartilage degeneration was severe (medial cartilage degeneration score for DMM 16+wks = 12.8±3.7), concordant with previous results (5–7). Four weeks after PMX, chondropathy was comparable to early DMM, but then progressed to severe cartilage degeneration with full thickness loss in some areas by week 12 (medial cartilage degeneration score for PMX 4+wks = 5.6±1.2, sham 4+wks = 0, PMX 12+wks

= 20.6±9.3, sham 12+wks = 0) (medial cartilage degeneration Suppl. Fig. 2D, total cartilage degeneration Suppl. Fig. 5L). One week after ACLR, disease severity was mild 1 week, both in the femur and tibia. By 4 weeks after ACLR injury, severe chondropathy with full thickness cartilage erosion was observed in the femoral condyles, while damage in the tibial plateau was mild (medial cartilage degeneration score for ACLR 1+wk = 3.85±1.8, sham 1+wk = 0.07±0.1, ACLR 4+wks = 16.4±11.2, sham 4+wks = 0) (medial cartilage degeneration Suppl. Fig. 2G, total cartilage degeneration Suppl. Fig. 5R). Naïve 26-week old mice showed no cartilage damage, while by 2 years of age, naïve mice showed mild joint damage in both knee joint compartments (for 26-week old mice: medial cartilage degeneration score = 0.4±0.55 and lateral cartilage degeneration score = 0.98±0.74, medial; for 2-yr-old mice: medial cartilage degeneration score = 4.72±1.19, lateral = 2.08±1.18), (medial cartilage degeneration Suppl. Fig. 2J, total cartilage degeneration Suppl. Fig. 5V), concordant with previously reported findings (8,9). Representative histological images of the medial side are shown in Suppl. Fig. 3. Representative images of the whole joint are shown in Suppl. Fig. 5, in addition to total joint cartilage degeneration scores and total joint synovial scores.

*Synovitis:* In addition to cartilage damage, we also assessed synovial hyperplasia, cellularity, and fibrosis for the four joint quadrants. Four and eight weeks after DMM, synovial changes were observed in the medial femoral and tibial compartments and were significantly more pronounced compared to age-matched shams (Suppl. Fig. 2B shows the total medial synovitis score, Suppl. Fig. 4A-C show the medial hyperplasia, cellularity, and fibrosis scores, Suppl. Fig. 5G shows total synovitis score). Synovitis scores were higher in the 4-week group compared to 8 and 16 weeks (Suppl. Fig. 2B). Significant increase in synovial cellularity was observed 4 weeks after DMM compared to the 8-week and 16-week timepoint (Suppl. Fig. 4B), while no difference was detected in fibrosis and hyperplasia of lining cells between the two timepoints (Suppl. Fig. 4A,C). Similarly, 4 and 12 weeks after PMX, the medial compartment showed pronounced synovitis compared to age-matched shams (Suppl. Fig. 2E shows the total medial synovitis score, Suppl. Fig. 4D-F show the medial hyperplasia, cellularity, and fibrosis scores, Suppl. Fig. 5M shows total synovitis score). The synovial changes peaked 4 weeks after surgery and went down by the 12-week timepoint (Suppl. Fig. 2E). Here too, increased cellularity was observed at 4 weeks compared to 12 weeks after PMX, while a trend of increased hyperplasia and fibrosis was observed at the early timepoint compared to the later timepoint (Suppl. Fig. 4D-F). Thus, findings in both surgical models are concordant with published literature reporting that synovitis scores were highest 4 weeks *post* DMM and reduced with time by week 12 (10).

Synovial changes were also detected 1 and 4 weeks after ACLR, compared to age-matched shams. No significant changes were detected between the 2 timepoints (Suppl. Fig. 2H). shows the total medial synovitis score, Suppl. Fig. 4G-I shows the medial hyperplasia, cellularity, and fibrosis scores, Suppl. Fig. 5S shows total synovitis score). These findings confirm our previous reports of robust synovitis in male mice 1 and 4 weeks after ACLR injury (4).

In naïve mice, 2-year-old mice showed mild medial and lateral synovial changes, a trend of increased cellularity and fibrosis was observed but was not significantly different compared to 26-week-old mice (Suppl. Fig. 2K) and (Suppl. Fig. 4K,L), confirming our recent findings (9). No difference was detected in hyperplasia of lining cells between young and old mice (Suppl. Fig. 4J) and (Suppl. Fig. 5W)

*Osteophytes:* Osteophyte width and maturity were assessed in the medial compartment for each model at both timepoints. DMM operated knees showed medium-sized osteophytes that mature with time over 16 weeks (average osteophyte width at 4 weeks= 183.9±76.6, at 8 weeks=

193.2±69.6, at 16 weeks=220.02± 46.39) (Suppl. Fig 2C). Similarly, four weeks after PMX, medium-sized osteophytes were observed, which further grew into large more mature osteophytes by week 12 (average osteophyte width at 4 weeks= 189.2±56.6, at 12 weeks= 314.7±72.7) (Suppl. Fig 2F). One week after ACLR, knees developed large chondrophytes at both the medial femoral condyle and tibial plateau, further growing and maturing into large, mature osteophytes 4 weeks after injury (average chondrophyte width at 1 week= 250.2±161.3, average osteophyte width at 4 weeks= 336.9.±106.5) (Suppl. Fig 2I). No osteophytes were observed in the lateral compartment of these models. Young naïve mice (26 weeks) did not show osteophytes, while older mice showed chondrophytes/small osteophytes in the medial compartment (average osteophyte width= 138.8±66.8) (Suppl. Fig 2L). Representative histological images of the medial joint are shown in (Suppl. Fig. 3). Representative histological images of whole joint, in addition to total joint cartilage degenerations scores and total joint synovial scores are shown in (Suppl. Fig. 5).

Representative histological images of the medial side are shown in Suppl. Fig. 3. Representative images of the whole joint are shown in Suppl. Fig. 5, in addition to total joint cartilage degeneration scores and total joint synovial scores.

**References**


1. Glasson SS, Blanchet TJ, Morris EA. The surgical destabilization of the medial meniscus (DMM) model of osteoarthritis in the 129/SvEv mouse. Osteoarthritis Cartilage. 2007 Sep;15(9):1061–9.

2. Knights CB, Gentry C, Bevan S. Partial medial meniscectomy produces osteoarthritis pain-related behaviour in female C57BL/6 mice. Pain. 2012 Feb;153(2):281–92.

3. Rzeczycki P, Rasner C, Lammlin L, Junginger L, Goldman S, Bergman R, et al. Cannabinoid receptor type 2 is upregulated in synovium following joint injury and mediates anti-inflammatory effects in synovial fibroblasts and macrophages. Osteoarthritis Cartilage. 2021 Dec;29(12):1720–31.

4. Bergman RF, Lammlin L, Junginger L, Farrell E, Goldman S, Darcy R, et al. Sexual dimorphism of the synovial transcriptome underpins greater PTOA disease severity in male

mice following joint injury [Internet]. Physiology; 2022 Dec [cited 2023 Jan 26]. Available from: http://biorxiv.org/lookup/doi/10.1101/2022.11.30.517736

5. Obeidat AM, Miller RE, Miller RJ, Malfait AM. The nociceptive innervation of the normal and osteoarthritic mouse knee. Osteoarthritis Cartilage. 2019 Nov;27(11):1669–79.

6. Little CB, Barai A, Burkhardt D, Smith SM, Fosang AJ, Werb Z, et al. Matrix metalloproteinase 13-deficient mice are resistant to osteoarthritic cartilage erosion but not chondrocyte hypertrophy or osteophyte development. Arthritis Rheum. 2009 Dec;60(12):3723–33.

7. Miller RE, Tran PB, Ishihara S, Larkin J, Malfait AM. Therapeutic effects of an anti-ADAMTS-5 antibody on joint damage and mechanical allodynia in a murine model of osteoarthritis. Osteoarthritis Cartilage. 2016 Feb;24(2):299–306.

8. Loeser RF. Aging processes and the development of osteoarthritis. Curr Opin Rheumatol. 2013 Jan;25(1):108–13.

9. Geraghty T, Obeidat AM, Ishihara S, Wood MJ, Li J, Lopes EBP, et al. Age-associated changes in knee osteoarthritis, pain-related behaviors, and dorsal root ganglia immunophenotyping of male and female mice. Arthritis Rheumatol. 2023 Apr 25;art.42530.

10. Shu CC, Zaki S, Ravi V, Schiavinato A, Smith MM, Little CB. The relationship between synovial inflammation, structural pathology, and pain in post-traumatic osteoarthritis: differential effect of stem cell and hyaluronan treatment. Arthritis Res Ther. 2020 Feb 14;22(1):29.
